# Robust and clinically relevant prediction of response to anti-cancer drugs via network integration of molecular profiles

**DOI:** 10.1101/301838

**Authors:** Marcela Franco, Ashwini Jeggari, Sylvain Peuget, Franziska Böttger, Galina Selivanova, Andrey Alexeyenko

**Affiliations:** Department of Microbiology, Tumor and Cell Biology (MTC), Karolinska Institutet, Stockholm, Sweden; Department of Cell and Molecular Biology, Karolinska Institutet, 171 77 Stockholm, Sweden; OncoProteomics Laboratory, Department of Medical Oncology, VU University Medical Center, 1081HV Amsterdam, The Netherlands; National Bioinformatics Infrastructure Sweden, Science for Life Laboratory, Box 1031, 17121, Solna, Sweden

## Abstract

In order to tackle heterogeneity of cancer samples and high data space dimensionality, we propose a method NEAmarker for finding sensitive and robust biomarkers at the pathway level. In this method, scores from network enrichment analysis transform the original space of altered genes into a lower-dimensional space of pathways, which is then correlated with phenotype variables. The analysis was first done on *in vitro* anti-cancer drug screen datasets and then on clinical data. In parallel, we tested a panel of state-of-the-art enrichment methods. In this comparison, our method proved superior in terms of 1) universal applicability to different data types with a possibility of cross-platform integration, 2) consistency of the discovered correlates between independent drug screens, and 3) ability to explain differential survival of treated patients. Our new in vitro screen validated performance of the discovered multivariate models. Finally, NEAmarker was the only method to discover predictors of both *in vitro* response and patient survival given administration of the same drug.

## INTRODUCTION

The problem known as the “dimensionality curse” [1],[2] - when a set of few (tens to hundreds) biomedical samples are described with a much larger number of molecular variables - undermines robustness of phenotype predictors. This was aggravated further when novel omics platforms expanded the variable space from thousands to nearly millions of potentially informative molecular features. In addition, profiling of cancer samples revealed that genomic alterations across tumors of the same type appear disparate and poorly overlapping [2]. As a result, variability between cancer samples is often higher than is assumed by the common parametric statistics [3]. Beyond a few success cases [4],[5] molecular cancer signatures have been hard to corroborate in a novel, independent cohort. Across a number of meta-analyses, conclusions about practical applicability of the signatures range from entirely negative [6],[7] to mixed or moderately positive [8]. The common understanding is that seemingly disparate individual events must be confluent to certain pathways that represent cancer hallmarks and pathways [9].

When, Modeling drug response *in vitro* was questioned by finding that molecular landscapes of cancer cell lines are be very different from those of original tumors [10]. A later, more comprehensive exploratory analysis demonstrated overall consistence of molecular aberrations between cell lines and primary tumors from matching cancer sites [9] – although these authors did not investigate the therapeutic relevance of discovered *in vitro* correlates. Haibe-Kains and co-authors published a discouraging comparison [11] between two large *in vitro* screens [12],[13]. After that conclusion and the following polemics [14], the urgent need in cross-platform and clinically based validation became even more apparent. It is dictated by both statistical and biological challenges, such as excessive data dimensionality, imperfect analytical tools, the heterogeneity of cancer genomes, and the downstream diversity of methylation and expression patterns [15]. Authors of one of the most up-to-date investigations still admitted that the ability of cancer cell drug screens to inform development of new patient-matched therapies…remains to be proven” [16]. On the clinical side, oncologists expected reports on patient-specific alterations in the light of knowledge available from computerized support systems [17]. In our view, these challenges could be most systematically addressed by summarizing sparse, disparate events at the pathway level via the global interaction network.

Adding omics data to clinical variables has demonstrated the potential for prediction of cancer disease outcome in a DREAM challenge [18]. One particularly winning strategy was to employ multigenic expression patterns. Such ‘meta-genes’ [19] were, despite the seemingly ‘network-free’ definition, nothing other than modules in a co-expression network, which allowed dimensionality reduction and a biological generalization. Another DREAM project revealed efficiency of summarizing gene expression in cancer cell lines over pathways [20].

Further, identifying patient sub-categories responsive to a treatment is more challenging than one-dimensional drug sensitivity or survival analyses. A practical method should profile individuals across the cohort, so that the profiles can be fit to clinical variables and covariates. Therefore, a crucial feature for biomarker discovery would be the ability to assign scores to individual samples rather than to derive feature-pathway associations from the whole data collection. In addition, further sample classification in a flow of new patients should not require re-running the analysis on the whole cohort, i.e. recalculating the data space, as is often the case.

In this work, we use acronym NEA to refer to a specific approach for network enrichment analysis, which ascends to the idea of accounting for the node degrees of individual genes [21]. Using that approach of significance estimation via comparing network connectivity to a null model, NEA [22],[23] can characterize experimental and clinical samples with pathway scores by accounting for sample-specific gene set relationships in the global gene interaction network. The pathway-level output is simple, uniform and statistically sound, so that it could be used in downstream analyses against arbitrary phenotype models. The ability to summarize rare alterations that cause the recurrent cancer phenotypes into pathway profiles provides higher statistical power, more information on the underlying biology, and robustness in phenotype prediction. However neither NEA nor alternative methods of pathway enrichment had been systematically applied to the task presented above: the discovery of biomarkers suitable for individual outcome prediction.

In the first section of Results, we provide a detailed explanation of the method NEAmarker and an instructive example, both in comparison with alternative methods. A representative set of such methods, was selected by investigating a wide range of earlier proposed algorithms and approaches. Since they were mostly designed for purposes different from ours, their applicability was often limited. In Methods (section “Alternative Methods of Pathway and/or Enrichment Analysis”), we discuss their principles, consider both applicability to biomarker discovery and software usability, and motivate our choice of methods presented in Figure 1 and Table 1. Thereby performance of our method is measured in parallel with using original gene profiles and those alternative enrichment methods: overrepresentation analysis (ORA), gene set enrichment analysis (GSEA, in two versions: [24], [25]), and signaling pathway impact analysis (SPIA, [26]). The outline and details of the comparative performance evaluation are reported in Results. More specifically, we: 1) assess content of relevant information in three published experimental *in vitro* drug screens [12] [13] [27] (dubbed CCLE, CGP, and CTD, respectively), 2) investigate preservation of this content across drug screens and then in one novel dataset, 3) perform a novel, small scale drug screen and demonstrate that the pathway-level multivariate models withstand the independent validation, and finally 4) validate the identified correlations in clinical treatment profiles from TCGA [28] (Table 2).

**Table 1.** Characteristic features of the alternative methods.

| Method | Type of input data | Allows data type integration | Level of input (samples) | Network analysis | Level of output (features) |
| --- | --- | --- | --- | --- | --- |
| original data | Any | - | All genes | - | *[same as input]* |
| ORA | Any | + | Altered gene sets | - | Functional gene sets |
| AGSEA | Expression | - | All genes | - | Functional gene sets |
| ZGSEA | Expression | - | All genes | - | Functional gene sets |
| SPIA | Expression | - | All genes | + | Functional gene sets |
| PWNEA | Any | + | Altered gene sets | + | Functional gene sets |
| GNEA | Any | + | Altered gene sets | + | Network gene nodes |

**Table 2.** Steps of analysis using alternative methods from Table 1.

| Step | What was evaluated | Figure | Scheme |
| --- | --- | --- | --- |
| 1 | Statistical power to detect correlates of drug sensitivity (fraction of significant correlates per dataset) | 3 | Within 3 published *in vitro* screens; within TCGA clinical datasets |
| 2 | Consistency of the discovered correlates between drug screens: cross-validation | 4 | Between 3 published *in vitro* screens |
| 3 | Consistency of multivariate models between drug screens: independent validation | 5 | From CTD *in vitro* screen to the novel ACT screen |
| 4 | Agreement between *in vitro* screens and clinical data | 6 | From 3 published *in vitro* screens to TCGA clinical datasets |

**Figure 1.**
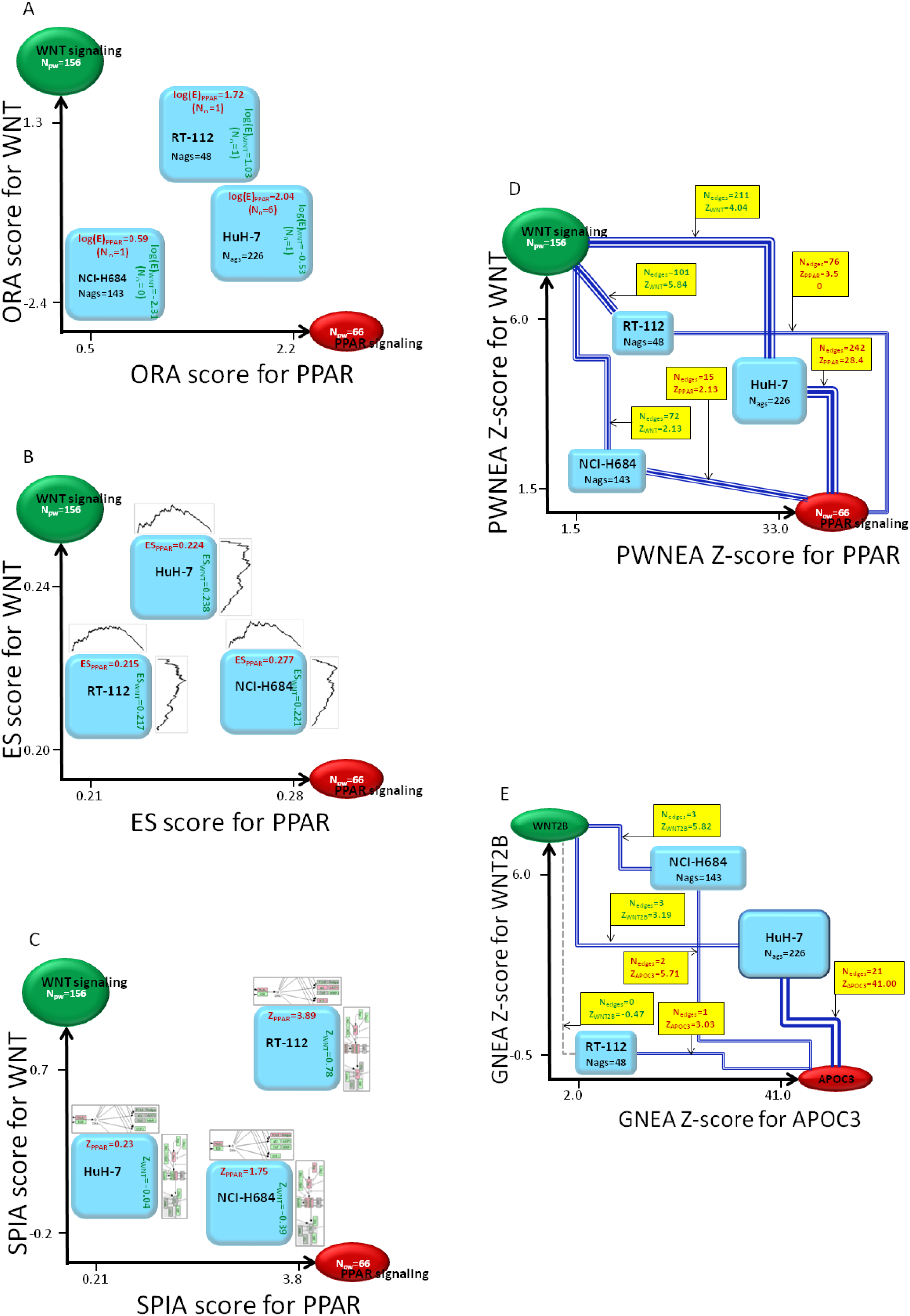
Rendering biological samples into pathway space with alternative enrichment methods. The placement of three cancer cell lines HuH-7, NCI-H684, and RT-112 in a 2-dimensional space of pathways ‘PPAR signaling’ and ‘WNT signaling’ (KEGG#03320 and KEGG#04310) (A, B, C, and D) or, alternatively, in a space of two key genes from these pathways (E) was done by using cell line-specific altered gene sets, AGS, which originated from transcriptomics data and contained 226, 143, and 48 member genes, respectively (AGS of class `significant.affymetrix_ccle`). A. ORA: enrichment of the three AGSs was analyzed against the two pathways (or, more generally, functional gene sets, FGS) using the overrepresentation analysis. The pathway enrichment scores were calculated from overlap between the gene sets. For clarity we here denote the pathway size *N*_*PW*_ which corresponds to *N*_*FGS*_ elsewhere in the article. Due to the relatively small gene sets sizes (*N*_*PW*_ and *N*_*AGS*_), a noticeable (*N*_*∩*_ *> 1*) and significant overlap was observed in just one out of six cases, which could limit the ORA sensitivity. B. GSEA was calculated using the full ranked gene lists from each cell line sample [24]. C. SPIA accounted for topological relationships of altered genes within the pathways. More weight was assigned to patterns of consistent up/down-regulation, i.e. where deregulated genes adjoined in regulatory cascades. Relatively disjoint regulatory events contributed with lower weights. The gene set submitted to SPIA can be of arbitrary size, up to full length, as in GSEA. The fold change values determine relative influence of the pathway genes. D. NEA: the coordinates of the three AGSs in the space of two pathways were determined via network enrichment analysis. The NEA z-scores (on the axes and in yellow boxes) were calculated via network connectivity rates between corresponding AGS and FGS by taking into account the numbers of AGS-FGS links (*N*_*edges*_ in yellow boxes) and the node topology of the member genes (Fig. 1 and Methods). The summarized connections between AGSs and FGSs are shown by blue compound edges that represent multiple individual gene-gene edges in the global network (*N*_*edges*_ ∼ line width). Individual edges within AGSs and within FGSs are not used in the analysis. E. GNEA: since the power of NEA to detect network enrichment was high, it was possible to apply NEA to the cell line AGSs versus individual gene network nodes WNT2B and APOC3 in the same way as it was done versus pathways in D. Even though the *N*_*edges*_ values were expectedly smaller than in B, four out of six Z-scores appeared rather high.

## RESULTS

### 1. Background

The main principle of NEA can be understood via comparison to gene set enrichment analysis in its simplest form, the so called overrepresentation analysis, ORA [29] (Fig. 1A). An experimental or clinical sample can be characterized by a set of altered genes (AGS), such as top ranking differentially expressed genes, or a set of somatic mutations, or a combination of these. The other component of the analysis is a collection of functional gene sets (FGS): pathways, ontology terms, or custom sets of biological importance. Importantly, FGS collections should summarize existing knowledge, being either expert curated or derived from experimental data. Enrichment scores of the AGSs can thus be used as the samples’ coordinates in the lower-dimensional FGS space. In ORA, enrichment is measured by the number of genes shared by the FGS and the AGS, given the sizes of the latter. NEA considers the network environment by counting the ***network edges that connect*** any genes of AGS with any genes of FGS (Fig. 1D). In both ORA and NEA significance can be evaluated with appropriate statistical tests. For NEA, this evaluation must be additionally normalized by topological properties of the network nodes. Due to the presence of different interaction mechanisms in the global network, NEA does not expect FGS genes to be altered themselves and therefore is capable of detecting enrichment of e.g. transcriptomics-based AGS in a pathway that operates by other mechanisms, such as trans-membrane signaling, phosphorylation etc. Compared to ORA, NEA holds other key advantages, such as exceptionally high power to detect enrichment in a global network, given the latter is sufficiently dense, i.e. when the median number of edges per gene is around 50. Hence, even smaller gene sets often connect to each other by multiple edges. An ultimately reduced FGS can even appear as an individual key network node. This gene-level network analysis, GNEA (Fig. 1E) provides a more focused alternative to the default analysis at the pathway level, PWNEA (Fig. 1D) and we therefore separately evaluated performance of PWNEA and GNEA in the present work.

The methods in Figure 1 implied different input, processing and output (Table 1). Accordingly, our data analysis procedure included the method-specific steps for sample/patient characterization, enrichment analysis, and phenotype modeling. In order to maximally adapt GSEA to our applications, we tested two different ways of ranking gene lists, AGSEA and ZGSEA (*Methods*) and present respective results separately. In sections 3…5 of Results, we report the results of systematic analyses of the experimental datasets with the alternative methods in order of increasing complexity (Table 2).

We begin by introducing an example of data analysis and interpretation (Fig. 2). Using data from the CGP in vitro screen, we observed a negative correlation between the PWNEA scores for pathway KEGG#00670 “One carbon pool by folate” for cell line AGS features `significant.affymetrix_ccle`, on the one hand, and sensitivity to methotrexate on the other hand (Spearman rank R = −0.248; p(H0) = 2.37e-06). The relatively low magnitude of the correlation is typical of such analyses and was explained by minor fractions of responders among all tested genotypes [14]. We compared cell lines which combined lowest sensitivity to methotrexate with highest PWNEA scores for KEGG#00670 (dubbed here Drug-/PW+) versus those possessing highest sensitivity and lowest PWNEA scores (Drug+/PW-) (ten cell lines in each set). Figure 2 (A,B,D,E) displays the network connectivity of the FGS KEGG#00670 “One carbon pool by folate” with AGSs for two cell lines (MPP89, ECGI10) of group Drug-/PW+ and two cell lines (RS411, A2780) of group Drug+/PW-. As an example, MPP89 obtained an NEA score of *Z*=8.09 (NEA FDR=4.3e-10; see details in Methods) because there were *n*_AGS-FGS_ = 19 edges in the network between its AGS and the FGS, against 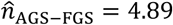 edges expected by chance. For comparison, the NEA *Z*-score for A2780 was as low as - 0.77 and insignificant. The negative sign indicated that the number of network edges *n*_AGS-FGS_ = 10 between the AGS and FGS was lower than the value expected by chance, 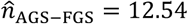. The expected numbers 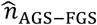 differed between MPP89 and A2780 due to the difference in the cumulative AGS degrees *N*_*AGS*_=2268 and *N*_*AGS*_=5823, respectively (shown in Fig. 2F). The high score for MPP89 (Fig. 2A) was likely influenced by the network node of formimidoyltransferase cyclodeaminase FTCD, which provided 14 out of the 19 edges. Although enrichment against the same FGS might have been enabled via entirely different AGS member genes, we note that it was not the case here: FTCD was a member of four out of the ten AGS of the group Drug-/PW+. Methotrexate is a cytostatic drug that inhibits dihydrofolate reductase, thereby blocking synthesis of tetrahydrofolate, the downstream production of folic acid, and finally that of thymidine. We can therefore hypothesize that overexpression of FTCD, an enzyme controlling the interconversion between formimidoyltetrahydrofolate and tetrahydrofolate [30], might have rescued the thymidine production by supplying extra tetrahydrofolate [31]. Since FDCD itself is a member of the “One carbon pool by folate”, the pathway could be, in principle, detected by another enrichment algorithm. But how have the alternative tested enrichment methods dealt with this pattern? Any noticeable correlations were absent. This might be explained by the fact that FDCD was the only consistently deregulated gene out of the whole pathway, which was a challenging situation for each of these methods. ORA is not well fit for cases of such an overlap (N=1). In GSEA, enrichment via a single highly ranked list member is usually not detectable. In its turn, SPIA could not gain enough statistical power in absence of consistent (adjoining) patterns of dysregulation in multiple genes. Finally, expression of FTCD itself did not significantly correlate with methotrexate sensitivity in CCLE and CGP transcriptomics datasets. More broadly, we did not find any genes of the “One carbon pool by folate” and the adjoining pathway KEGG#00790 “Folate biosynthesis” which would significantly (by requiring q-value <0.05) correlate with methotrexate sensitivity at either gene expression or somatic mutation levels.

**Figure 2.**
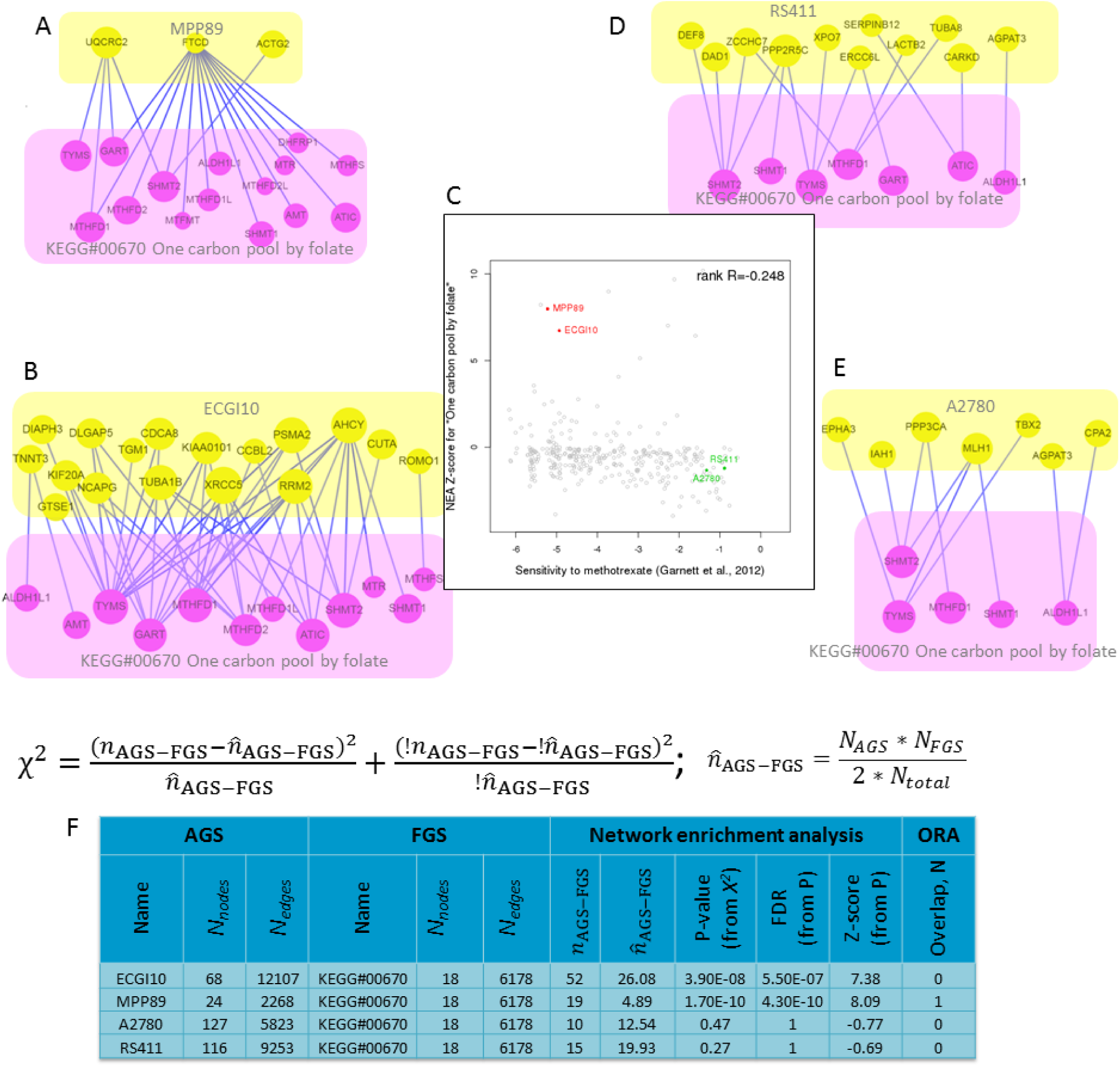
Network enrichment analysis of four cell line AGSs with differential response to methotrexate. While using AGSs of class `significant.affymetrix_ccle`, the response of cancer cell lines to methotrexate in CGP screen correlated with NEA scores (pane F) in regard to FGS “One carbon pool by folate” (pane C). The methotrexate-resistant cell lines MPP89 and ECGI10 (panes A and B) received higher NEA scores since the numbers of edges *n*_AGS-FGS_ connecting them to the FGS significantly exceeded those expected by chance, 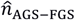 (52 vs 26.02 and 19 vs. 4.89, respectively; pane F). For comparison, the sensitive lines RS411 and A2780 (panes D and E) had fewer edges than expected (15 vs 19.93 and 10 vs. 12.54, respectively) and therefore received lower, negative scores. The table in F and the sub-networks in A, B, D, and E were created via the web-site for interactive NEA https://www.evinet.org/.

For comparison, AGS of the other resistant cell line, ECGI10, did not share any genes with the target pathway (Fig. 2B), although still received a higher NEA score. In this case, the summarized connectivity was not dominated by a single network node of the AGS or of the FGS. Here the drug resistance could potentially have been mediated by the DNA repair protein XRCC5 or by the adenosylhomocysteine hydrolase AHCY, which were earlier reported to be implicated in methotrexate resistance [32] and folate metabolism [33], respectively. Unlike the upregulated FTCD in MPP89, both these genes were strongly downregulated in ECGI10. This emphasizes another feature of NEA: genes may be included into AGS due to alterations in an arbitrary direction, i.e. both over- and under-expression, hyper- and hypo-methylation, increased and decreased copy number etc. Therefore higher and, respectively, lower NEA scores cannot be traditionally interpreted as activation or suppression of the given pathway (FGS) but rather indicate a general ‘pathway perturbation’. Hence the pathway “One carbon pool by folate” was unperturbed in the low-scoring cell lines A2780 and RS411, i.e. the latter did not exhibit features that could connect specifically to the pathway.

### 2. Construction of sample-specific AGS

In order to analyze data from the *in vitro* cancer cell screens and the primary tumor samples (TCGA) in the same manner, we constructed AGSs by following the same platform-specific approaches. Intuitively, having an AGS that is too big or too small could deteriorate specificity or sensitivity of NEA. Therefore, in order to prove that differences are not due to selecting AGS genes in a specific way, we tested and compared a number of options for AGS compilation. Mutation-based AGSs were created by first listing all point-mutated genes in each given sample (which might include hundreds and even thousands of passenger mutations) and then retaining only those with significant network enrichment against the rest of the set. This approach [34] had been proposed for distinguishing between driver and passenger mutations - hence the filtering should reduce noise by enriching AGSs in driver genes. Next, AGSs from gene copy number and expression data included genes most deviating from the cohort means. This was achieved by using one of the three alternative algorithms (see Methods). Again, even such deviant gene sets could still be too large, e.g. due to listing copy number-alterations over extended chromosomal regions. In order to compact these, alternative AGS versions were derived by retaining only genes with significant network enrichment for signaling and cancer pathways or for the mutation-based AGS of the same sample, which reduced the AGS lists 3-10 fold. An alternative to using gene copy number data would be to account for respective mRNA expression levels. While this approach is subject of ongoing discussion, we have observed [34] that many known copy number drivers did not exhibit this correlation and therefore we decided not to filter copy number data by the gene expression feature. Finally and as an extra option, we merged platform-specific AGSs into combined AGSs.

### 3. Statistical power to detect correlates of drug sensitivity

The goal of this first, exploratory analysis was to compare the different methods and feature classes in their ability to explain the differential drug sensitivity. To this end, we counted features significantly associated with a phenotype after adjusting the respective p-values for multiple testing. For example (Suppl. Fig. 1), we analyzed associations between point mutation profiles of cancer cell lines [35] and cell lines’ sensitivity to each of the 203 anti-cancer drugs from Basu et al. [27]. The fraction of low p-values (e.g. *p(H_0_)* < 0.001) in the total number of statistical tests did not exceed that expected in absence of any associations and therefore no genes received q-values (adjusted p-values) [36] below 0.05. On the contrary, the correlation analysis of gene expression [12] against the same drug sensitivity profiles discovered nearly 15,000 patterns of association between gene expression and drug sensitivity (out of in total 18,900 x 203 = 3,836,700 tests) with *p(H_0_)* < 0.001. After the adjustment, more than 2500 of these gene-drug pairs remained significant at *q* < 0.001. These two examples demonstrate how dramatically the information content could vary depending on the feature type and data origin.

Applying this approach to the *in vitro* drug screen data, we evaluated features of different types and classes. Respectively, in TCGA data we measured correlations of features with survival of patients who received one of the 42 frequently used drugs in any of the eight cohorts. We systematically compared different feature types, i.e. original data from high-throughput platforms and NEA scores as well as classes within the types (e.g. transcriptomics data from Affymetrix vs. Agilent vs. RNA sequencing). We also analyzed the relative performance of different AGS classes.

Overall, the NEA scores at both pathway level (PWNEA) and individual gene node level (GNEA) contained either approximately the same or larger amounts of information on drug sensitivity compared to the original gene profiles (Fig. 3). In the drug screen data analysis, the ORA, PWNEA, and GNEA features performed apparently better than the respective original point mutation, gene copy number, and gene expression data. In the TCGA data analysis, the advantage of PWNEA and GNEA over both ORA and original gene profiles for particular drugs was even more pronounced, although not always overall significant. Among the platforms for the *in vitro* screens, Affymetrix data by far outperformed mutation data, copy number, and combined AGSs. In TCGA datasets, RNA sequencing performed better than Affymetrix (the former data was not available for the cell lines). In general, transcriptomics datasets much more frequently manifested correlations with drug sensitivity than gene mutations and copy number datasets (Suppl. Fig. 3). While this observation is not new [9], the most obvious explanation should be that most of the genome alterations were insufficiently frequent for the statistical tests. As an example, less than 10% of the genes in the BRCA cohort had point mutations in more than 1% of the tumors and therefore the analysis did not gain enough statistical power. mRNA expression profiles were, on the contrary, available for most of the genes. We also assessed relative performance of the different AGS classes. From each dataset with continuous values we created AGSs of fixed size (`top.200` and top.400) as well as sets of variable size where genes were included based on significance as referred to the cohort mean (`significant`) and, in addition to the latter, tested for network enrichment toward cancer gene sets (`significant.filtered.mini`) or any signaling pathways (`significant.filtered.maxi`). As illustrated in Supplementary Figure 3, the different classes yielded variable results. We evaluated consistency and significance of these differences using the same Kolmogorov-Smirnov test as in Figure 3 on the gene copy number and expression datasets for cell lines and TCGA samples (Suppl. Table 1). This evaluation, however, did not lead to an unequivocal conclusion. In the cell line datasets, the fixed size AGSs performed significantly better, while in the TCGA datasets the situation was rather opposite.

**Figure 3.**
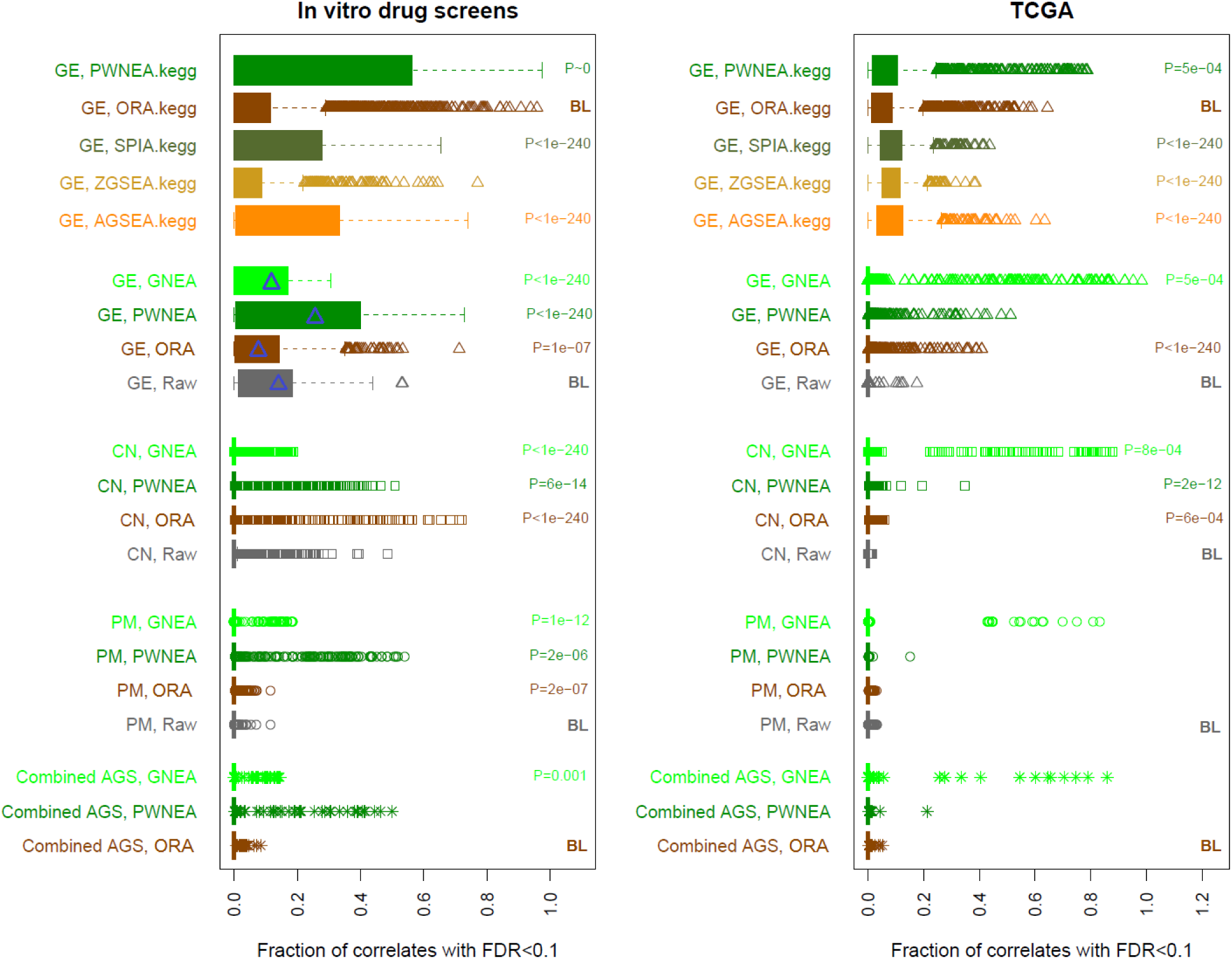
Comparison of the potential performance of different features, methods, and data types. The top 5 boxplot rows (labeled “.kegg”) present results obtained using the limited set of 197 KEGG pathways using gene expression data (label “GE”). Next, since ORA, PWNEA, and GNEA did not require intra-pathway topology and could accept any data type, the rest of boxplots present tests on the full set of 328 FGS (the respective PWNEA and ORA values for GE might differ, since the KGML gene sets were somewhat different from the core KEGG version). Each boxplot element combines correlation values between either features for a given class (labeled at the vertical axis) and for either the *in vitro* response to drugs in the three screens (left pane; in total 365 tests of 320 distinct drugs) or for the survival of patients who had been treated with drugs (right pane; 42 drugs in the eight TCGA cohorts). As an example, we calculated Spearman rank correlations between sensitivity of cell lines to drug RITA and transcriptomics features of these cell lines: either original Affymetrix (CCLE) gene expression profiles (18900 genes) or enrichment profiles of cell line specific AGSs of class `top.400.affymetirx_ccle` produced by GSEA (328 FGS features), pathway-level NEA (PWNEA; the same 328 features), and gene-level NEA (GNEA, in which 19027 nodes in the global network were treated as single-gene FGSs). The p-values of Spearman correlations between features and the drug sensitivity were then adjusted for multiple testing. The fractions of adjusted p-values below 0.1 became X-coordinates for the plot. The four examples (indicated by the blue markers), respectively, gave fractions 1837/18900=0.097; 23/328=0.070; 78/328=0.236; and 2090/19027=0.110. Each boxplot element combined such fraction values for each drug from each screen or TCGA cohort as well as all alternative AGSs classes for ORA, PWNEA, and GNEA. The features are grouped by type of profiling (original data, ORA, PWNEA, and GNEA as grey, brown, dark green and bright green, respectively) and by data type (point mutations, PM; copy number alterations, CN; and gene expression, GE). The p-value is shown when a category produced significantly (p<0.001 by Kolmogorov-Smirnov test) more non-zero patterns (i.e. fractions with FDR<0.1) than the respective baseline category (labeled “BL”). The boxes contain data points within 25-75th percentile intervals (i.e. between quartiles Q1 and Q3). The maximal whisker length, MWL, is defined as 1.5 times the Q1-Q3 interquartile range (i.e. the box length). Whiskers can extend to either the MWL or the maximal available data point when the latter is below MWL. Markers thus correspond to data points that extend off the box by more than the MWL value.

While the original features manifested considerable correlations in a number of classes, fractions of significant correlations were largely inferior when compared to NEA classes. In general, the different methods could be ranked by potential sensitivity in the following order: original gene profiles < [either ORA or ZGSEA] < [either AGSEA or SPIA]< [either PWNEA or GNEA]. However even upon adjustment for multiple testing, we did not draw ultimate conclusions from significance of these correlations. This exploratory analysis only informed us on the relative Type II error rates (i.e. sensitivity, or statistical power to detect correlation), suggesting that multiple alternative methods and data types were potentially predictive of drug sensitivity. In order to evaluate robustness of these predictions we proceeded to the validation step as described below. We also note that only ORA, PWNEA, and GNEA could provide means for integrating omics data from different platform (by simple merging of AGS lists), where ORA was apparently inferior.

### 4. Consistency of the discovered correlates in different drug screens

In order to test reproducibility of the drug-feature associations in alternative experimental settings, we used data from three *in vitro* drug screens: CCLE [12], CGP [13], and CTD [27]. A comparison between CCLE and CGP screens was earlier presented in [11]. The CTD drug screen was published later and provided additional shared compounds for our cross-screen analysis (31 in addition to the 16 available to Haibe-Kains and colleagues). Similarly to these authors, we found that the association values between drug sensitivity and original features only weakly agreed between the drug screens.

Albeit weak, these correlates were still significantly concordant across screens. Fig. 4A presents examples of between-screen rank correlations when using original gene expression profiles, ORA, PWNEA, and GNEA features. When comparing results from screens by CGP [13] and CTD [27], the correlation values between Affymetrix expression data and sensitivity to navitoclax ranged from R=0.31 (original gene profiles) to R=0.81 (GNEA). More systematic analyses demonstrated (Fig. 4, B and C) that using AGS features in PWNEA and GNEA considerably strengthened the concordance compared to the original gene profiles and AGSs in ORA. For example, by requiring across-screen rank correlations above 0.6, four NEA feature classes based on gene copy number performed better than any original copy number class. Under the same rank correlation threshold, eight out of ten transcriptomics NEA classes and all those based on point mutations were superior to the respective original data classes. Results obtained with ORA were, again, inferior to those from NEA and the summarized ranking appeared as: [original gene profiles and ORA] < PWNEA < GNEA. In the tests using 197 KGML-KEGG pathways and gene expression data, SPIA and AGSEA were somewhat superior over PWNEA.

**Figure 4.**
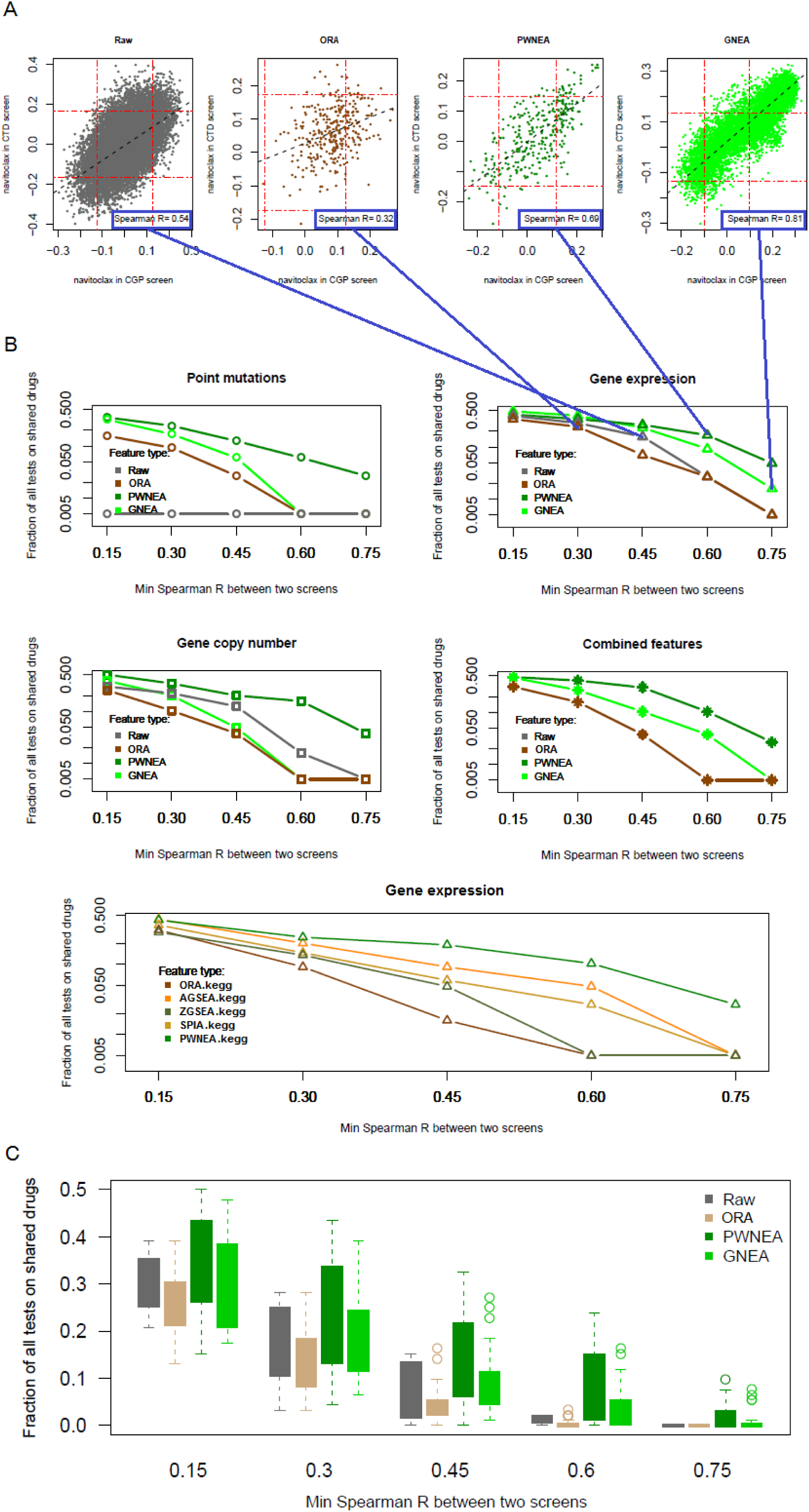
Consistency of drug-feature associations between drug screens. For each drug shared by any two of the three *in vitro* drug screens (in total 47 cases), we calculated rank correlation between drug-feature rank correlation coefficients in the two screens. A. Agreement of drug-feature rank correlation coefficients between CGP and CTD screens of sensitivity to navitoclax using Affymetrix data as original gene expression values Affymetrix_CCLE (left pane) and AGS features of class `significant.affymetrix_ccle` profiled with ORA, PWNEA, and GNEA (other panes). The agreement in this case was worst upon using ORA profiles (rank R=0.32), whereas GNEA profiles performed best (rank R=0.81). The red lines indicate the levels of false discovery rate (the correlation p-value adjusted by Benjamini-Hochberg) FDR=0.1. The grey diagonal line is the linear regression fit. B. Fractions of cases with rank correlation value above each of the five specified thresholds on example AGS classes. The features are grouped by type of profiling and by data type identically to Fig. 3. Four example values from A are mapped to the gene expression plot in B. In order to characterize sensitivity to each of the 47 shared drugs, we used here, in parallel with respective original gene profiles, AGS features of one class of each type: significant.filtered.exome.mini (PM), significant.filtered.snp6.mini (CN), and `significant.affymetrix_ccle` (GE), and significant.filtered.combined.maxi (‘combined’). The advantage of GNEA (with the exception of “Point mutations” and “Gene copy number”) and PWNEA became apparent at the highest cutoffs R>0.60 and R>0.75. C. Similarly to B, fractions of values above each of the five specified thresholds were calculated for *all* classes and combined for all data types. For certain AGS classes, PWNEA and GNEA produced correlates highly conserved across screens (R>0.6) in as many as 5-10% of cases.

We validated drug sensitivity profiles of four anti-cancer compounds, tested previously in the CTD screen - RITA, PRIMA-1^MET^/Apr-246, nutlin and JQ1 - in a new *in vitro* screen, named ACT. The activities of these compounds were re-tested in a panel of 20 cancer cell lines (the ACT set) for which the CCLE gene expression and point mutation profiles data were available. The wide response ranges indicated sufficient differential response across the ACT set. Similarly to the results in Fig. 3, both original gene profiles and NEA features showed significant, moderately strong correlation with drug sensitivity, which demonstrated the potential of multivariate models for drug sensitivity prediction.

As shown above, the original gene profiles were poorly preserved across drug screens. Therefore, we compared the CTD results with those from ACT screen in a more relevant multivariate approach using the elastic net method [37]. Starting from all available features, each model was finally reduced to a much smaller subset. Multi-variate models are notoriously prone to over-fitting when the number of variables exceeds the number of samples. For this reason, validation on independent sets has become an essential requirement in such studies [38]. We thus created CTD-based models using cell lines not found in the ACT screen. The comparison was also streamlined by using only the data from CCLE Affymetrix and point mutation datasets versus two respective feature AGS classes `mutations.mgs` and `significant.affymetrix_ccle`. Using other classes produced similar results (data not shown). Figure 5 demonstrates that by applying the same set of elastic net parameters, in every case it was possible to obtain a descriptive model from CTD drug screen data with a number (4…129) of non-zero terms and then substantiate the model (possibly with a poorer performance) using the ACT data in a smaller cell line set. For each modeled case, we compared observed and predicted drug sensitivity values. The most important observation was that in all instances without exception the signs of these correlations were consistent between CTD model and ACT validation, i.e. negative correlations in the training set remained negative upon validation.

**Figure 5.**
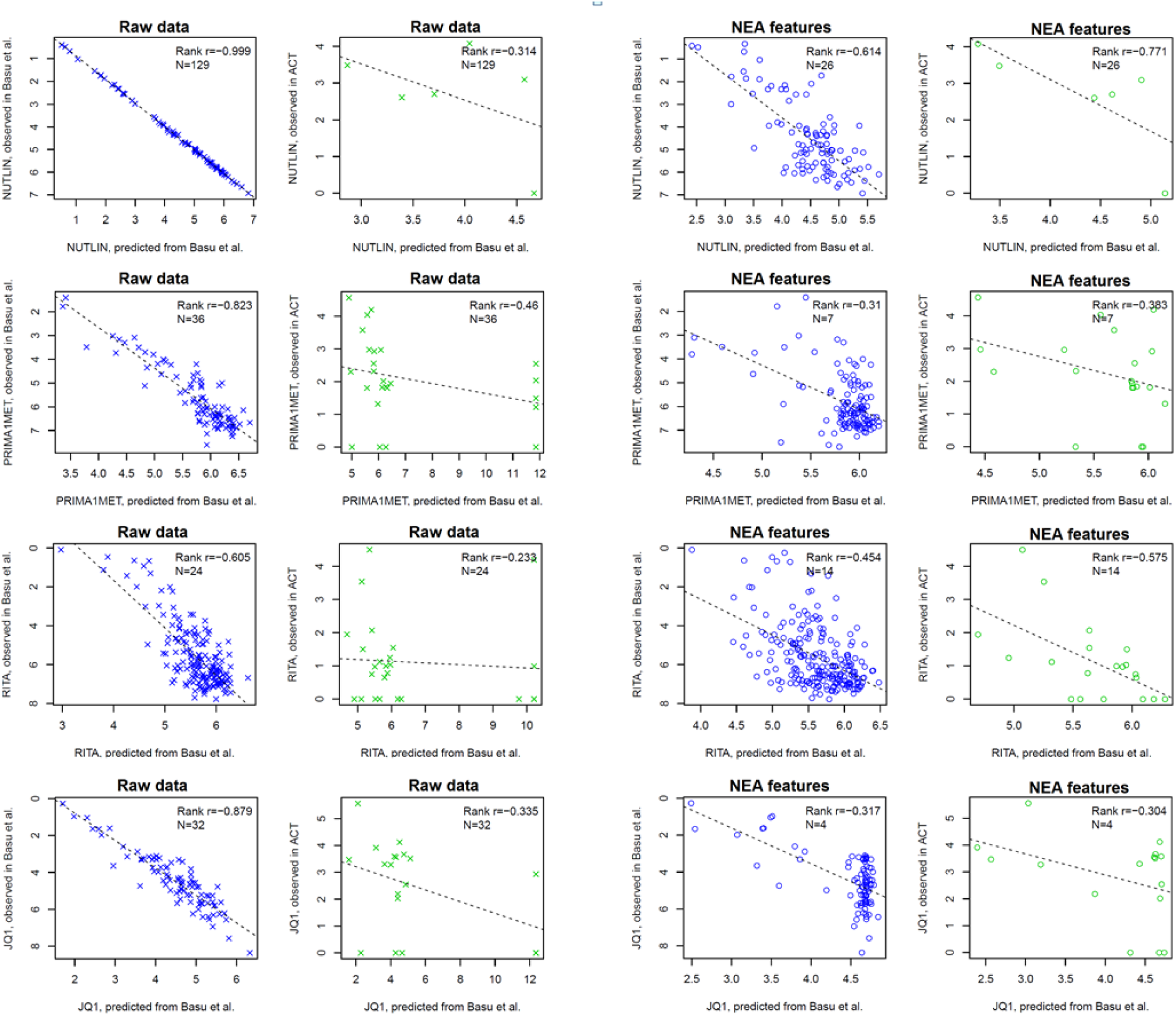
Predicted versus observed drug sensitivity across cancer cell lines in discovery versus validation screens. The predictive models for four compounds tested in the published CTD screen were validated in our ACT screen. Elastic net models were built under multiple cross-validation inside the training set (columns 1 and 3, blue) and then tested on non-overlapping sets of cell lines of the ACT screen (columns 2 and 4, green). Input variables were either original gene point mutation and expression profiles (columns 1 and 2, crosses) or PWNEA scores derived from these datasets for each cell line (columns 3 and 4, circles). Legends in each plot display the values of Spearman rank correlation between observed and predicted values (’Rank r’) and the number of non-zero terms in the model (’N’). Parameter `alpha` for the shown plots was set to 0.9. Since the drug sensitivity values from CTD screen were inverted compared to the other screens, the correlations are presented as negative values (see also inverted vertical scales for “observed in Basu et al.”). Detailed plots for models built under different `alpha` parameters are found in Supplementary Files glmnetModels.Basu_vs_new.raw.pdf and glmnetModels.Basu_vs_new.pwnea.pdf.

Overall, the performance of the original profile models on the validation sets appeared comparable to that of PWNEA. However importantly, the former had much more freedom in model term selection since the initial feature space was around two orders of magnitude larger than that in PWNEA. Consequently, despite the rigorous cross-validation and feature selection implemented in the `glmnet` algorithm, using the original profiles generated more complex models (see the number of terms per model, N) which fit the training sets better (and were clearly overfitting in the case of nutlin). At the validation step however, the performance of the models based on original data significantly worsened - whereas the performance of PWNEA-based models remained at roughly the same level (all results obtained under variable parameters can be found in Supplementary Files glmnetModels.Basu_vs_new.raw.pdf and glmnetModels.Basu_vs_new.pwnea.pdf). This result essentially corroborated the previous conclusion about lower robustness of the original gene profiles compared to NEA.

### 5. Agreement between *in vitro* screens and clinical data

A more challenging task was to identify a conservation of associated features between the *in vitro* drug screens and clinical application of the same drugs. Any trustworthy setup of such an analysis would be very complex, so that even cross-validation and adjustment for multiple testing could not guarantee unbiased probabilistic estimation. Thus, the final judgment should had been made after a biologically independent *ad hoc* validation from the *in vitro* to the clinical domain. Even though the TCGA collection did not provide correctly balanced, randomized cohorts for estimation of relative risks, error rates etc., our task was simplified by only needing to compare the methods’ performance. In the eight largest TCGA cohorts, we counted how many significant *in vitro*-detected features correlated with survival of patients who received same drug [28], (https://tcga-data.nci.nih.gov/docs/publications/tcga/; Suppl. Table 4), [39]. More specifically, molecular features of each class that were significantly correlated with sensitivity to a drug in cell lines were required to be also significantly correlated with patient survival in a TCGA cohort. Our survival analysis accounted for clinical covariates available from TCGA (Suppl. Table 4), which enabled estimating the ‘net’ effects of molecular features.

We matched correlates of same data types in CCLE and TCGA (although possibly obtained using different omics platforms, e.g. Affymetrix microarray from CCLE could be matched to RNA-seq from TCGA etc.). Then we determined whether correlation p-values of individual features, in their turn, correlated between *in vitro* and TCGA data, i.e. if genes or FGSs with high (respectively low) correlation with drug response *in vitro* tended to correlate in the same manner with the patients’ response. Due to the testing of alternative AGS classes, respective numbers of matching pairs in ORA, PWNEA, or GNEA were an order of magnitude higher than in raw data (column 2 in Table 3). Therefore we coupled this calculation with a significance test by randomly permuting feature and sample labels. Altogether, the permutation tests indicated that point mutation and copy number data had zero true discovery rates (TDR), i.e. their correlation p-values were preserved not more than expected by chance (see column 3 in Table 3). On the contrary, the TDR levels were substantial (0.02…0.805) for gene expression data and for AGSs processed with each of the enrichment analyses.

**Table 3.** Conservation of drug sensitivity correlates between the *in vitro* drug screens and clinical applications.

| Feature type | | No. of available "feature X drug" tests | True discovery rate by permutation test, $p(H_0) < 0.01$ | No. of usable correlates, so that $p(H_0) < 0.001$ and rank $R > 0.2$ | | | | | | | | |
| --- | --- | --- | --- | --- | --- | --- | --- | --- | --- | --- | --- | --- |
|  |  |  |  | All TCGA cohorts<br>(% of available correlates) | Bladder carcinoma,<br>BLCA | Breast carcinoma,<br>BRCA | Colon adenocarcinoma,<br>COAD | Glioblastoma multiforme, GBM | Lung adenocarcinoma,<br>LUAD | Lung squamous carcinoma , LUSC | Ovarian carcinoma,<br>OV | Prostate adenocarcinoma,<br>PRAD |
| Original gene profiles | Point mutations, PM | 360 | 0 | 0 | 0 | 0 | 0 | 0 | 0 | 0 | 0 | 0 |
|  | Copy number alterations, CN | 522 | 0 | 0 | 0 | 0 | 0 | 0 | 0 | 0 | 0 | 0 |
|  | Gene expression, GE | 1080 | 0.149 | 0 | 0 | 0 | 0 | 0 | 0 | 0 | 0 | 0 |
| Enrichment analysis | ORA | 9014 | 0.033 | 78<br>(0.8%) | 0 | 56 | 0 | 3 | 3 | 8 | 8 | 0 |
|  | PWNEA | 8822 | 0.146 | 305<br>(3.5%) | 18 | 60 | 20 | 52 | 51 | 45 | 59 | 0 |
|  | GNEA | 8630 | 0.805 | 505<br>(5.9%) | 15 | 84 | 46 | 113 | 93 | 72 | 82 | 0 |
| Enrichment analysis,<br>GE on KGML KEGG only | ORA | 5252 | 0.025 | 21<br>(0.3%) | 0 | 5 | 0 | 2 | 4 | 2 | 8 | 0 |
|  | ZGSEA | 1080 | 0.037 | 8<br>(0.7%) | 0 | 2 | 0 | 0 | 1 | 0 | 5 | 0 |
|  | AGSEA | 1080 | 0.020 | 7<br>(0.6%) | 0 | 7 | 0 | 0 | 0 | 0 | 0 | 0 |
|  | SPIA | 1080 | 0.048 | 3<br>(0.3%) | 0 | 0 | 0 | 3 | 0 | 0 | 0 | 0 |
|  | PWNEA | 4988 | 0.241 | 364<br>(7.2%) | 44 | 83 | 20 | 79 | 27 | 19 | 92 | 0 |

At the next step (remaining columns of Table 3) we calculated the numbers of significant cases that would also be practically usable, i.e. had both lower p-values (<0.001) and rank correlation values higher than 0.2. No such cases were identified in the gene expression data. ORA, PWNEA, and GNEA yielded 0.8%, 3.5%, and 5.9% of practically usable cases, respectively. Interestingly, most (56 out of 78) of the ORA cases were identified in the breast cancer cohort, whereas the preserved PWNEA and GNEA correlations distributed uniformly across all the TCGA cohorts, except prostate cancer which cohort shared only one drug with one *in vitro* screen. Remarkably, the separate test using the 197 KGML KEGG pathways also demonstrated superiority of PWNEA over ORA, ZGSEA, AGSEA, and SPIA - despite the reasonably good performance of the latter two in the *in vitro* analyses presented above. Thus at this crucial validation stage, robustness of the data types while translating drug sensitivity correlates between *in vitro* and clinical applications increased in the following order: [point mutations and gene copy number changes] < [gene expression] < [ORA, ZGSEA, AGSEA, and SPIA] < PWNEA < GNEA.

The purpose of this analysis was to prove systematically significance of the produced correlates, and we reiterate that using the original data did not seem efficient: although many transcriptomics profiles correlated with drug sensitivity, those patterns could not be traced back to the *in vitro* screens.

Most of the consistent NEA features were obtained for AGS based on gene expression data (Suppl. Table 2). They were identified for docetaxel, gemcitabine, and paclitaxel in BRCA (see the cancer cohort notation in Table 3); for dexamethasone, erlotinib, and topotecan in GBM; for gemcitabine in LUAD; and for gemcitabine, paclitaxel, tamoxifen, and topotecan in OV. While using gene copy number data, consistent PWNEA and GNEA features were found only for GBM (dexamethasone and topotecan). Consistent features that correlated with the response to cisplatin (LUSC) belonged to the combined, multi-platform types. Only one consistent GNEA feature was based on somatic mutation analysis (gemcitabine in LUSC), although it did not match all the criteria. Below we present promising features found in the drug screens, which were also predictive of survival if the drug was administered in a TCGA cohort.

The cancer emergence and progression were earlier linked to tissue inflammation through the NOD-like receptor signaling [40]. We found that the corresponding pathway score correlated with survival in ovarian carcinoma patients treated with topotecan (Fig. 6A).

**Figure 6.**
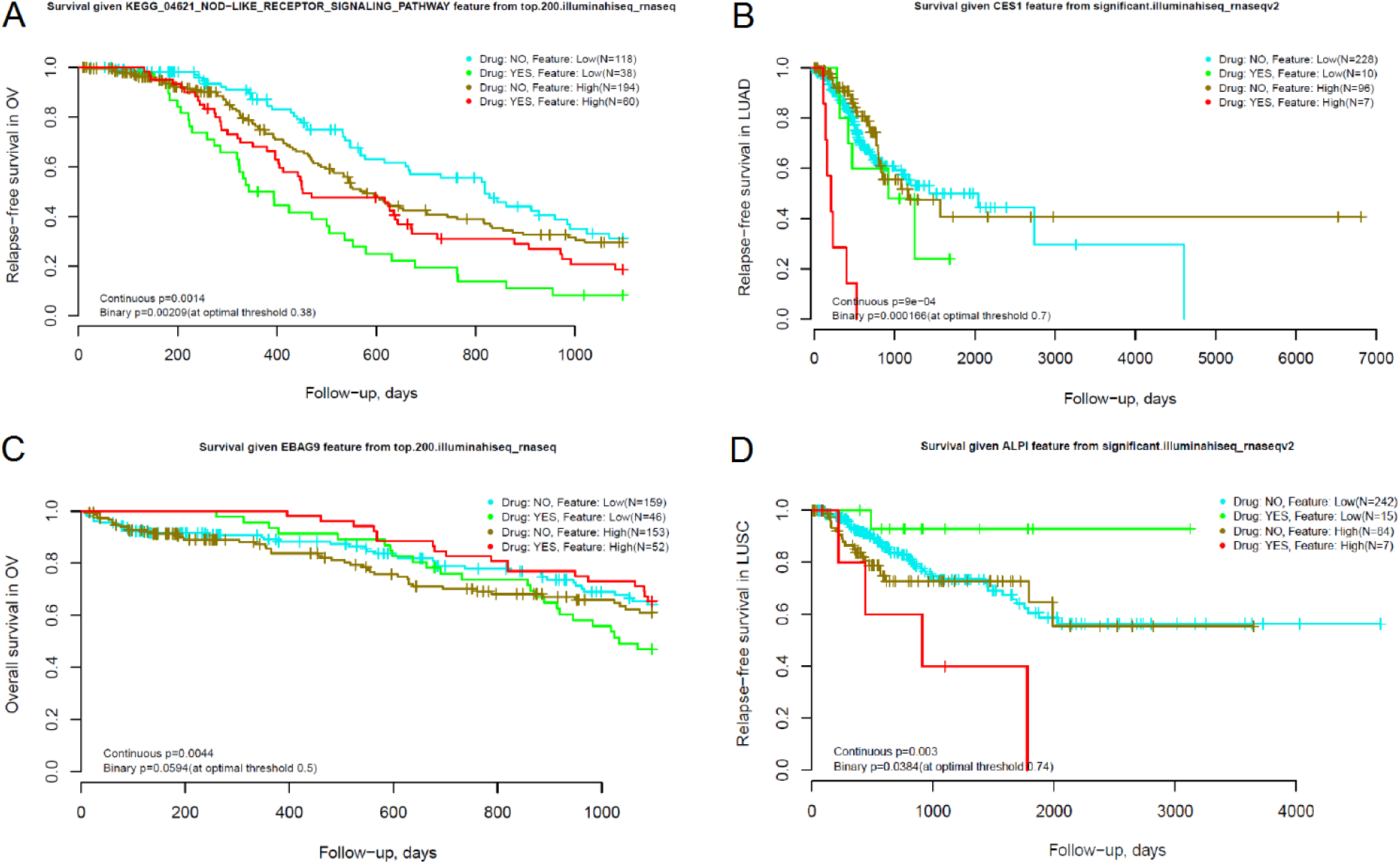
Clinical performance of NEA features discovered in drug screens. Each TCGA cohort was split into four categories by two factors: 1) administration of the specific drug and 2) predictive feature value (pathway or individual gene score, indicated in the plot header), each above and below a threshold. The primary feature evaluation employed p-values calculated in the continuous score space, i.e. without splitting the patient cohort into binary classes by factor (2). Then the binary classifications by the both factors were used for visualization (as “treated/untreated” for the drug and at the quantile “optimal threshold” value for the quantitative NEA feature). The p-values are shown for the both alternatives. The plots present differential survival upon treatment with topotecan in ovarian carcinoma (A and C), gemcitabine in lung adenocarcinoma (B), and vinorelbine in lung squamous cell carcinoma (D).

Carboxylesterases (CESs) are capable of hydrolyzing gemcitabine [41] - for instance, CES2 slows down hydrolysis of the gemcitabine pro-drug LY2334737 [42]. We identified as many as 31 gene-wise NEA features which correlated with relapse-free survival in lung adenocarcinomas treated with gemcitabine. This list of network nodes from GNEA included CES1 (Fig. 6B), CES2, CES7, and a number of cytochromes with possible involvement in the catabolism of xenobiotics. Many of these genes were AGS members in both the gemcitabine-sensitive cell lines and in patients who responded to the gemcitabine treatment and – at the same time - they themselves were members of KEGG pathways 00980 “Metabolism of xenobiotics by cytochrome p450”, 00983 Drug metabolism – other enzymes, and 00982 “Drug metabolism – cytochrome p450”. Consequently, the ORA and PWNEA analyses detected enrichment of these pathways in the same patients. However the pathway scores correlated with response to gemcitabine neither in the CCLE and CTD screens nor in the LUAD cohort) and therefore would be useless as biomarkers. The gene expression profiles of carboxylesterases and cytochromes in cell lines and primary tumors did not correlate with gemcitabine response either. EBAG9 had been implicated previously in ovarian cancer progression [43], but it has not been shown to affect response to topotecan. Indeed, in the datasets of our study the expression of the gene itself correlated neither with cell line sensitivity to topotecan nor with patient survival. However, the GNEA features for EBAG9 as a network node did correlate with sensitivity to topotecan *in vitro* (`top.200.affymetrix_ccle`; p(H_0_)=4.2×10^−11^) and with overall survival of OV patients (Fig. 6C) (`top.200.illuminahiseq_rnaseq`; p(H_0_)=4.4×10^−4^ during the 3-year follow-up time while accounting for clinical stage as a covariate).

The intestinal-type alkaline phosphatase ALPI is known to be a modulator of cancer cell differentiation [44] and cytoprotection [45], [46]. In our analysis its GNEA feature was, in parallel with eleven others, negatively correlated with sensitivity to vinorelbine *in vitro* (`gnea.significant.affymetrix1`; p(H_0_)=1.1×10^−07^) and with overall survival of OV patients (p(H_0_)=0.003; Fig. 6D).

This setup could not eliminate possible confounding effects from multi-drug treatment history and clinical factors that might determine administration of specific drugs. Nonetheless, the NEA scores apparently explained the differential sensitivity to anti-cancer drugs in a much more robust and efficient manner than the original data.

A visual inspection of the survival curves in Fig. 6 sheds light on usefulness of these tentative biomarkers in a clinical setting. As an example, in a 1-year survival perspective, relative risks (RR) would either increase (Fig. 6A,C) or decrease (Fig. 6B,D) given higher NEA scores of the patient samples. By using this fixed follow-up interval and the cohorts of limited size, the confidence intervals at the 95% level would be rather broad: ln(RR) = 0.405 (95% CI: [-0.07…0.88]); ln(RR) = −2.061 (95% CI: [-3.99…-0.13]); ln(RR) = 2.211 (95% CI: [-0.70…5.12]); ln(RR) = −2.181 (95% [CI: −5.15. 0.78]) for Fig. 6A…D, respectively. The fractions of patients who might benefit from using these predictors could be estimated in terms of absolute risk reduction as 0.17, 0.62, 0.08, and 0.25. Inversely, the “number needed to treat”, i.e. how many patients should be treated for one individual to benefit from the new test would have been 6.00, 1.60, 12.91, and 3.94, respectively [47]. However, additional responders could be detected by using other markers, used in parallel. As an example, beyond the “NOD-like receptor signaling pathway” at Fig. 6A, the response to topotecan in ovarian cancers similarly correlated with KEGG pathways “One carbon pool by folate” and “Bacterial invasion of epithelial cells” as well as with the GO term “Cytokine activity” (not shown). Predictions made with these markers would overlap only partially and therefore can complement each other. We presume that such discoveries should ultimately be evaluated by independent validation and careful clinical development. In fact, our combined analysis of independent cell screen and clinical results gave a first example of such validation.

## DISCUSSION

We have presented a way of using network enrichment scores for prediction of drug response and demonstrated its advantage compared to the conventional analyses of original gene profiles and alternative enrichment methods. In comparison to the latter, the NEA scores correlated stronger with drug sensitivity and were preserved better between independent screens. Multivariate models using NEA scores proved more compact and, at the same time, robust when re-tested on newly obtained data. Finally, corroborating *in vitro* phenotypes in corresponding clinical applications was possible by using the method NEAmarker but not by original profiles or alternative methods.

In our view, the advantages of our approach are due to the following features of network-based data interpretation: 1) combining major types of molecular interactions in a biologically relevant way, 2) summarizing seemingly disparate molecular alterations at the level of pathways and processes, and 3) enabling lower-dimensional statistical analysis. In addition, network views provide better grounds for biological interpretation and mechanistic studies. The types of evidence behind the edges (such as protein-protein interactions, mRNA co-expression, sub-cellular co-localization) might contribute to the integrated network differently. We refer to the previously published comparative analyses of the contributions [48], [34], [49]. The poor performance of the individual gene analysis and ORA could be explained by the excessive dimensionality of the former and poorer sensitivity of the latter (Fig. 1A). In addition, the ability to use smaller and hence more specific AGS could have provided extra advantage of NEA over ORA and GSEA. On the other hand, NEA could also deteriorate on AGS of insufficient size when using sparser networks (around 10^4^…10^5^ edges) and networks with many missing nodes. These potential limitations were established earlier [34] and we tried to avoid them in the present work by using e.g. the dense network from data integration. Also there could be no edges connecting an AGS to a specific FGS (even though such cases would still have certain variability of NEA scores due to variable values of 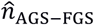. We admit that a future, more comprehensive version of NEA might adopt advantages of the alternative enrichment methods by employing full gene lists (as in GSEA) and intra-pathway topology (as in SPIA). Indeed, at two steps of our analysis these methods demonstrated performance comparable to that of NEA (Fig. 3 and 4).

A common problem of method benchmarking is the unavailability of ground truth. In our case, too, we did not possess a set of truly existing molecular correlates of drug sensitivity. Comparing alternatives by the total number (fraction) of positives would not enable a proper control of the false positive rate. In similar situations, when it was impossible to distinguish between true and false positives, authors often chose to present biologically sensible examples, such as enrichment of a pathway pertinent to the problem [22], [50] or correlation with a known drug target [15]. In the present work, we evaluated concordance of phenotype correlations between different, independently collected datasets. This allowed us to circumvent the problem of false positives via a more compelling prove: the methods were compared by the fractions of corroborated findings, which would be extremely unlikely by chance.

We started with analysis of drug screens using samples from The Cancer Cell Line Encyclopedia [12] profiled for somatic point mutations, gene copy number changes, and gene transcription [35],[13],[12]. Consequently, in TCGA cohorts we focused on the same data types. The individual molecular phenotypes were characterized with AGSs compiled using a number of alternative methods. The analysis provided a primary comparison of their relative performance but – at the current stage – did not enable definite conclusions about performance of the different AGS classes. Indeed, AGS of fixed size (`top.N`) versus variable size (`significant`) compared differently in the cell lines versus the TCGA data (Suppl. Table 1). Further in the analysis of consistency *in vitro* versus clinical results, these classes were almost equally represented (Suppl. Table 2). We have also seen differences between different filtering approaches in AGSs of classes `significant.mini` and `significant.maxi` (Suppl. Fig. 2). Therefore an issue to be investigated further is the comparative performance and robustness of different feature classes, platforms etc. Importantly, multiple platforms’ data can be integrated into combined AGSs. Although in our analysis such AGSs did not perform much better than platform-specific ones (most likely due to the domination of transcriptomics data), a more detailed evaluation should be done, including new platforms from TCGA and elsewhere, such as DNA methylation, protein phosphorylation etc. Given the diversity of carcinogenesis routes and the multiplicity of respective molecular mechanisms, combining platforms appears essential and most promising. Incorporation of approaches from sparse linear regression modeling, SPIA, GSEA, and PARADIGM certainly represent promising ways in this direction.

The statistical power of NEA was obviously far from full. As an example, there were 13 drugs for which the numbers of tested cell lines and patients treated in TCGA cohorts were sufficient for a significant estimation. For four drugs out of these 13, no reliable correlates could be found. One instructive example could be irinotecan, prescribed to 25 and 22 patients in COAD and GBM cohorts, respectively. The interesting feature of irinotecan is that its pharmacokinetic pathway involves the same enzymes as that of gemcitabine (Fig. 6B), namely CES1, CES2, CYP3A4, CYP3A5 and some others (https://en.wikipedia.org/wiki/Irinotecan#Interactive_pathway_map) – although the enzymes here work in an opposite direction: they activate irinotecan rather than degrade as they do to gemcitabine. Nonetheless, relevant GNEA scores might have been informative for response to irinotecan. The patients’ response was sufficiently differential, too: while all the irinotecan-treated patients relapsed, the time to relapse varied from 78 to 1265 days. However, we did not observe almost any sensible correlation of the pathway genes neither as GNEA features nor as raw gene expression profiles. In the GNEA framework, this elucidated a lack of network linkage between the AGSs of responders (or non-responders) to the irinotecan pathway.

Further, our FGSs were created by third-party sources and never meant to be used in NEA. Thus, another step for NEA-based biomarker discovery would be the compilation of novel, specifically optimized FGSs. Ultimately, one could compile *de novo* pathways - similarly to the approach by [51] - but specifically informative of the drug response or disease prognosis. An example of such a functional set could be the presented above combination of the ten carboxylesterases and cytochromes.

Finally, given the low overlap of member genes between individual AGS, it is important to establish how AGS-level biomarker panels would practically summarize gene-level information and organize the accompanying statistical framework. Ways to compile and employ multi-platform AGSs, optimal FGS design, and construction of NEA-based biomarker panels should therefore become the topics of future studies.

## MATERIALS AND METHODS

### Drug screens

#### Cell lines used in ACT screen

In this analysis, we used 20 cancer cell lines for which molecular data could be found in the CCLE Affymetrix set as well as in both CCLE and COSMIC point mutation sets: A375, HCT116, HDLM2, HT29, JVM2, K562, L428, MCF7, MDAMB231, MV411, NB4, PL21, RAJI, RKO, SJSA1, SKBR3, SKNAS, SW480, T47D, and U2OS. Eight of these cell lines had also been included in the CTD screen (A375, HCT116, HT29, MCF7, PL21, RKO, SW480, and U2OS). In order to avoid overlap in the multivariate models, we excluded these eight cell lines while training the original models from the CTD data and only used them in the validation set.

#### Assay for cell proliferation used in ACT screen

Cell proliferation was estimated with the WST-1 assay (water soluble tetrazolium). Briefly, cells were incubated with each drug for 72 hours in a 96-well plate. At the end of this period, they were incubated with WST-1 reagent (Roche) for 2 hours. Absorbance at 450nm was measured following the instructions from the manufacturer. The cell proliferation rate compared to that in the control was calculated.

For adherent cultures, cells were attached overnight before adding the compounds. For hematological malignancies, the compounds were added simultaneously with seeding cells. The initial cell density was chosen so as to avoid confluence at the end of the assay. Each compound was applied in six consecutive 3-fold dilutions. In all cases except JQ1, the stock for each drug was established at the concentration based on efficacy determined individually for each drug. Final concentrations were for RITA: 0.01, 0.04, 0.12, 0.37, 1.11, 3.33 µM; for Apr-246/PRIMA-1-met: 0.3, 1, 3, 9, 28, 83 µM; for Nutlin-3a: 0.14, 0.41, 1.23, 3.7, 11.11,33.33 µM. For JQ1 the cell lines HDLM2, HT29, MCF7, RAJI, RKO, SJSA1, SKBR3, and SW480 were tested using the final concentration range 1.66. 0.007 µM in 1:3 serial dilutions. However later we found it necessary to raise the concentration by one order of magnitude, so that the final concentrations for the rest of the cell lines were 16.66, 5.55, 1.85, 0.61, 0.20 µM. Then we respectively adjusted IC50 values for the first group as if they were tested under the final concentrations. This was done by incrementing the initial-stock IC50 values of HDLM2, HT29, RKO, and SW480 by *log_3_*(10) ≈ 2.09. The cell lines MCF7, RAJI, SJSA1, SKBR3 did not show any sensitivity while using the initial stock (IC50 = 0), so that their IC50 values upon JQ1 treatment were declared missing.

IC50 was defined as the drug concentration inducing a 50% reduction in cell proliferation compared to the control. In the quantitative analysis, we used a universal scale for all the four drugs where units 1…6 stood for dilution steps (1=1:300; 2=1:900; 3=1:2700; 4=1:8100; 5=1:24300 and 6=1:72900). Sensitivity to compounds was expressed in IC50 values varying from 0 (insensitive to compound) to 6 (fully sensitive to compound).

IC50 values and p-values of the model parameters were calculated using function drm from R package drc [46]. The model form (argument fct) was chosen as LL.4, where model parameters Lowest and Highest were fixed at cell proliferation rates 0% and 100%, respectively, while parameters slope and IC50 were left unfixed.

The IC50 values are provided as Supplementary File IC50values.ACTscreen.xlsx.

#### CCLE screen

Barretina et al. [12]analyzed cell line sensitivity to 24 drugs in 504 cell lines. These authors considered a range of numeric sensitivity metrics for their analysis and finally preferred ‘normalized activity areas’. These original units were calculated as areas under compound response curves where higher values corresponded to higher sensitivity so that 0 stood for ‘insensitive to compound’ and 8 corresponded to ‘full sensitivity’.

Further, the activity area values were normalized for unequal luminescence in the assay. We rendered them normally distributed by log-transformation. Thus the values in our analysis range from −3.00 meaning ‘insensitive to compound’ to +2.31 meaning ‘maximal sensitivity’.

#### CGP screen

Garnett et al. (2012) [13] analyzed 138 drugs in 714 cell lines. They used a combination of IC50 and the slope parameter to achieve the most complete description of responses. We decided to use the AUC as a single feature that reflects the both values. AUC was originally provided in the same table and ranged from 0% (fully sensitive) to 100% (insensitive). To approach the normal distribution, we transformed the values as *log*(1 - AUC), so that now they ranged from −8.11 meaning ‘insensitive to compound’ to 0 meaning ‘maximal sensitivity’.

#### CTD screen

The authors (Basu et al., 2013) [27] mainly used areas under curve (AUC) for their quantitative analysis of 203 drugs in 242 cell lines. We reproduced this approach in our study. In completely insensitive cases, the full area under eight experimental points reached 8, whereas 0 stood for full sensitivity. Thus, the scale of this screen was inverted compared to the other screens, which was considered in all calculations.

### Molecular data

#### Gene expression

The profiling was performed in CCLE study using Affymetrix GeneChip^®^ Human Genome U133 Plus 2.0 Array and in CGP study by Affymetrix GeneChip^®^ HT Human Genome U133 Array plate. The expression datasets were normalized as described by the authors and made public. Expression profiles in the CTD study were from CCLE. It has been shown earlier [11] that disagreement between CGP and CCLE could be attributed to the usage of different transcriptomics datasets only to a minor extent. We checked both the CCLE and CGP expression profiles and concluded that the latter provided poorer statistical power in regard to drug sensitivity as well as lower coverage of both genes (13891 unique mapped gene symbols vs. 18900 in CCLE) and cell lines (622 vs. 1034). For these reasons, we used the CCLE dataset in all the presented analyses. Expression values *x* of the downloaded datasets were transformed to *log_2_*(x).

### Gene Copy Number

CCLE, CGP, and CTD all employed Affymetrix SNP 6.0 microarrays for gene copy number detection. We downloaded the CCLE dataset [12] for 994 cell lines. In addition, we downloaded COSMIC data [35] independently produced by the same platform and then post-processed in three different ways to provide total, absolute copy number per gene, number of copies of the minor allele, and a binary classification of gene copy number values into gain vs. loss. All datasets were used as downloaded, without further processing or normalization.

### Point mutations

CCLE provided point mutation data on sequencing of 1667 genes in 904 cell lines. In addition, we downloaded COSMIC data from exome sequencing of 1023 cell line genomes, which mapped to 19759 gene symbols. Mutation data from the both screens were used in the binary form, i.e. all specifying attributes were neglected.

Following the same approach, we employed TCGA data on somatic point mutations reported in MAF files. The column ‘Variant_Classification’ contained a number (more than 15) different codes, most frequent being Missense_Mutation, Nonsense_Mutation, and Silent. Since the latter constituted around 25% of the total number of somatic mutations reported in the eight cancers - which would not significantly affect the false positive and true discovery rates - we analyzed records with any such codes as potentially associate ed with drug response.

### Alternative Methods of Pathway and/or Enrichment Analysis

We evaluated a number of existing multivariate, enrichment-based, and/or network analysis methods that could be potentially useful in the proposed analysis, accounting for their complexity, applicability to different experimental designs, and the ability to analyze individual samples rather than the whole cohort. Various statistical algorithms have been proposed to quantify functional relevance of pathways and other gene sets by accounting for gene network topology.

A number of methods can generate sparse regression models via network-based regularization, i.e. account for topological relations between potential predictors (typically gene expression variables). The regularization is based on certain assumptions, such as that e.g. term coefficients of neighbor nodes should be zeroes or non-zeroes simultaneously [52], that edge confidence weights should influence penalties on the model coefficients [53], or that there exists equivalence (or at least parallelism) between connectivity of nodes and covariance of model terms [54], [55]. Advanced regularization of linear models in these methods often demonstrated promising efficiency [56]. However being very sophisticated, these models proved hard to tailor to novel, specific experimental designs. Notably, it was not feasible to include additional covariates or interaction terms which would be necessary for e.g. analyses similar to the one described in the present work not even in the dedicated survival analysis method DegreeCox [57]. Using pathway membership information for summarizing cross-pathway linkage was proposed in [58] - however, adjusting its error rate model to other purposes has not been straightforward.

Technically, individual scores that estimate samples’ uniqueness as compared to the rest of the collection can be obtained already from ORA, i.e. from the simplest analysis of dichotomous 2×2 tables applied to sample-specific gene sets [59], [60], [61], [62], also called “class I” in the classification by Huang et al.[29]. For comparison, the most popular gene set enrichment analysis, GSEA [24] has been usually applied to finding pathway enrichment in gene lists pre-ranked by cohort-wise statistics. As an example, Haibe-Kains et al. [11] analyzed correlations between drug sensitivity and molecular features calculated on whole *in vitro* drug screens [12],[13] which are among the datasets re-analyzed in this article. Those pathway enrichment scores represented correlates of drug sensitivity over the whole screened collections rather than characterized individual cell lines. Likewise, Iuliano and co-authors [63] matched molecular landscapes to survival in cancer sample cohorts in order to reverse-engineer relevant pathway and network structures. Thus, global methods often employ powerful, heavily optimized statistical techniques and are used for sample exploration or differential expression analysis [27], [64] but cannot serve features for phenotype prediction in novel cell lines or tumors. An overview of network applications in cancer studies [65] showed that, indeed, most of the existing methods enabled exploratory analyses, discovery of driver genes and pathways as well as splitting a cohort into molecular subtypes, but did not characterize individual cases.

A number of hybrid approaches, such as SPIA [26] and iPAS [66] were also capable of calculating sample-specific pathway scores. However, their scores were based on gene expression values, which excluded the using of other data types. A genuinely integrative multi-omics method PARADIGM [67] (the program is currently distributed only via a company web portal), on the contrary, accounted for combinations of events in the chain DNA->mRNA->protein activity. As input, it required well characterized regulatory relationships – a complete set of which would rarely be available. Also, similarly to the former group of methods, it relied on comparing cancer to normal samples. Those dramatic alterations between the normal and cancer tissues encompassing thousands of genes would mask more fine-grained features that determine between-tumor heterogeneity, differences between sensitive and refractory cases etc. This requirement also precluded analyzing data where normal matches are missing, such as the widely used in our analysis cancer cell lines. Finally, EnrichNet [50] has been an algorithm closest in spirit to NEA: by using random walk with restart (hence not limited to 1-step network distances), it can trace AGS-FGS relationships via network paths. However it existed only in a single-AGS, web-based implementation and therefore was also not available for testing it the present analysis.

Even though individual enrichment scores can be correlated with phenotypes, they have still been rarely used in predictor models. In the case of ORA and GSEA, the major reason was that the enrichment is mostly detectable for large FGSs (hundreds to thousands genes), but such are unlikely to characterize functional differences between tumors - while compact, specific, and discriminatory gene sets tend to escape their limits of statistical power. Nonetheless, Drier et al. [68] have explored cancer cohorts with pathway-level sample scores derived from gene expression data in a quantitative way and found that certain sample clusters can be associated with patient survival. On the other hand, the network-based methods have been developed only recently and are therefore ‘too young’ to have been exploited fully. Above, we have also mentioned the network-based regularization of multiple regression models where inclusion of gene terms into the models was essentially coupled to their co-expression.

We finally decided to include in our testing, in parallel with PWNEA (pathway level NEA) and GNEA (gene node level NEA), the following methods:

1. Using original gene profiles from respective omics platforms;
2. ORA, over-representation analysis which was capable of working on exactly the same AGS and FGS as PWNEA;
3. GSEA on full ranked gene lists, applying two alternative methods:
  a. AGSEA, ranking by absolute gene expression value,
  b. ZGSEA, ranking by deviation of gene expression from the cohort mean;
4. SPIA, measuring the pathway perturbation via known intra-pathway topology.

Using GSEA and SPIA was restricted to only transcriptomics data. SPIA, in addition, could only be run on pathways with known topology, which limited the set of available FGS to 197 KEGG pathways available in KGML format. This created an additional, specific line of testing on a limited collection of input data and FGSs for the methods ORA, AGSEA, ZGSEA, SPIA, and PWNEA (see Fig. 3, 4 and Table 3).

### Network Enrichment Analysis (NEA, PWNEA, and GNEA)

#### Network

The network was based on the FunCoup method [48] with consecutive merging of five more resources as described and benchmarked previously [34]. The results of that benchmark indicated that FunCoup was superior to STRING (a method similar to FunCoup in terms of scale and the size of input data collection, [69]), mostly due to the latter broadly using prokaryotic evidence and therefore less specific in cancer-related analyses. The second conclusion from the benchmark was that adding to the FunCoup network edges of curated databases significantly improved its performance. We therefore added the FunCoup-based network with functional links from KEGG [70], CORUM [71], and PhosphoSite [72], MSigDB transcription factor-related part, [73]), and an own reverse-engineered network [34]. The resulting network thus combined a wide range of molecular mechanisms, functional relations, and metrics from high-throughput data sets: physical protein-protein interactions, membership in same protein complex, membership in the same pathway, correlation of mRNA profiles, correlation of protein abundance values, protein phosphorylation, coherence of GO annotations, concordance of upstream regulators (transcription factors and miRNAs), co-localization in same sub-cellular compartments, similarity of phylogenetic profiles etc. It contained 974,427 edges (links) between 19027 nodes (distinct human gene symbols).

#### Altered gene sets, AGS

Point mutation data (mutation gene sets):

- mutations.mgs: point-mutated genes that proved to be significantly NEA-enriched to either KEGG pathway set #05200 Pathways in cancer or to the full set of point-mutated genes annotated in the given genome (the approach described by Merid et al. [34]).

Gene copy number and expression data:

- top.200 and top.400: genes with copy number or mRNA expression value that in the given genome was among top 200 or top 400 most deviating from the gene’s cohort mean using the one-sample Z-score. Each AGS thus had a fixed size, regardless of formal significance.
- significant: most deviating from the gene’s cohort mean (same as above), but selected only if below the formal significance threshold (Benjamini-Hochberg [74] adjusted p-value<0.05). These AGSs had variable sizes, depending on the significance criterion.
- significant.filtered.mini: members of the respective significant set had, in addition, to be also significantly NEA-enriched to either KEGG set #05200 Pathways in cancer or to mutations.mgs set of the same sample (whichever NEA score passed the significance threshold NEA FDR=0.05).
- significant.filtered.maxi: members of the respective significant set were required to be significantly NEA-enriched to any of the signaling pathways (including all cancer ones) or to mutations.mgs set of the same sample.

Combined (multi-platform) AGS:

- significant.filtered.combined.mini: a merge of all sets of type significant.filtered.mini.
- significant.filtered.combined.maxi: a merge of all sets of type significant.filtered.maxi.

For convenience, AGS labels refer also to the platforms and sources, e.g. `top.200.cn_ccle`, `significant.filtered.maxi.affymetrix_ccle` etc.

### FGS

The functional gene sets, FGSs, were AGS counterparts in the analysis. The main collection of 328 FGS was based on the KEGG pathways, the full collection of which was complemented with a number of separately published cancer pathways as well as specific GO terms corresponding to cancer-relevant signaling or hallmarks of cancer (around 70 cancer- and signaling-related gene sets from Reactome, Gene Ontology, WikiPathways and literature). Another approach was applied to enable compatibility with GSEA and SPIA. These methods were designed and are most suitable for analyzing expression data and, apart from that, SPIA was applicable only to pathways with well characterized intra-pathway topology. We therefore employed a special set of 197 KEGG pathways for which the topology was available in KGML files and tested on it SPIA, GSEA, ORA, and PWNEA exclusively gene expression data (these results were separately labeled as ORA.kegg, SPIA.kegg, AGSEA.kegg, ZGSEA.kegg, and PWNEA.kegg). The analysis on the FGS collection is referred to as pathway-level NEA (PWNEA).

In the other version of our analysis, called gene-wise NEA (GNEA), we treated each of the 19027 network nodes, regardless of their pathway or GO annotation, as a single-gene FGS.

### Method

The major principles of NEA were described earlier [22]. In the current implementation, we evaluated enrichment of AGS versus FGS by the formula:

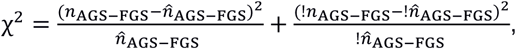

where !*n* means “complement to *n*”, i.e. all global network edges that did not belong to *N*_*AGS-FGS*_. The number of links expected under true null, i.e. by chance, was determined by:

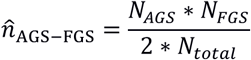

Node connectivity values (numbers of all edges for each given node) were pre-calculated by the algorithm in advance, given the input network. Then *N*_*AGS*_ and *N*_*FGS*_ reported the sums of connectivities of member nodes of AGS and FGS, respectively, and *N_total_* was the number of edges in the whole network. Since it was desirable to provide normally distributed values for the downstream analyses (linear modeling, correlation, survival), we calculated p-values from the *X*^2^ statistic *p*(*H_0_*)=*f*(*X*^2^) using function pchisq available in R language and then re-calculated corresponding *z*-scores from the p-values as *Z=F(p*(*H_0_*)) with function qnorm. Since *X*^2^ is only defined on the non-negative domain, the *z*-scores were coerced negative in cases of depletion, i.e. when

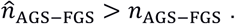

An important feature of GNEA (gene-wise NEA) is that its enrichment estimates are, on average, based on fewer network edges compared to PWNEA, so that often *n*_AGS-FGS_ =0. In such cases, the enrichment score is negative and the difference 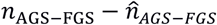 reduces to 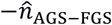, which, in its turn, is a function of cumulative connectivity values *N*_*AGS*_ and *N*_*FGS*_. In other words, lower NEA scores are then assigned to AGS-FGS pairs with more highly connected member nodes.

The steps of NEA described above can be performed with functions available in R package NEArender (https://cran.r-project.org/web/packages/NEArender/).

### Signaling pathway impact analysis, SPIA

The method by Tarca et al. [26] was implemented as an R package `SPIA`. The authors presented it as combination of two p-values: `pNDE` from common analysis of overrepresentation of differentially expressed genes in KEGG pathways and `pPERT` from a perturbation analysis by accounting for topological relations of the same genes within each KEGG pathway. Since the authors claimed that `pNDE` values are no different from p-values from the trivial ORA, we used the pure `pPERT` values from function spia (while the performance of ORA was evaluated separately). In order to get normally distributed values for our analyses, `pPERT` were transformed to *Z*-scores and signed according to the SPIA “Activated/inhibited” status as:
`Z.spia=qnorm(pPERT/2, lower.tail=F)*ifelse(s1$Status=="Activated", 1, −1);`

### Gene Set Enrichment Analysis, GSEA

The R implementation of GSEA was downloaded from http://software.broadinstitute.org/gsea/msigdb/download_file.jsp?filePath=/resources/software/GSEA-P-R.1.0.zip (see also https://software.broadinstitute.org/cancer/software/gsea/wiki/index.php/R-GSEA_Readme). While GSEA possesses a sophisticated toolbox for significance estimation via permutation tests, we needed only the enrichment score and therefore calculated only the core ES values via function `GSEA.EnrichmentScore`. Normally, GSEA has been used for analyzing gene rankings from multi-sample analyses with replicates, such as a *t*-test of an experimental versus control group. The single-sample GSEA (so called ssGSEA) needed for our analysis was described by Barbie et al. [25]. They produced sample-specific lists by ranking genes by absolute expression values in each given sample. We implemented this analysis under acronym AGSEA. However this approach might miss sample specificity. As an example, such ubiquitously expressed genes as GAPDH, RPS16, and RPS11 were found among the top 10 items in more than 90% of the CCLE cell line transcriptomes. For this reason, we additionally implemented and tested ranking genes in each sample by z-scores, i.e. by the standardized deviations from the genes’ means across the whole cohort. Using this option, dubbed ZGSEA, was similar to mode `topnorm` for calculating AGS in function `samples2ags` of our package `NEArender`.

### Overrepresentation analysis, ORA

The overrepresentation analysis, ORA estimated the significance of overlap between AGS and FGS in 2×2 tables. We did it via Fisher’s exact test using the function `gsea.render` in the R package `NEArender` described above. In order to get ORA values normally distributed, the `“estimate”` values from function `fisher.test` were augmented with a pseudo-score 0.1 and log-transformed.

### Correlation between drug sensitivity and molecular features

In each of the four drug screens, we quantified correlation between the cell line sensitivity to each drug and each of the molecular features *F* according to a general model of the form:

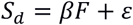

where *ε* denotes residual, i.e. unexplained by feature *F*, variance. The features were either original gene profiles from the three platforms (point mutations screens, copy number arrays, and expression microarrays) or scores from GSEA, or scores from the two NEA modes, PWNEA and GNEA, i.e. pathway-level network enrichment scores and single-gene network enrichment scores, respectively. All data sets, except the point mutation set, contained continuous variables and were thus analyzed using Spearman rank correlation. The point mutation data were analyzed using a one-way ANOVA model with two levels of *F*: any mutation versus wild type. P-values of both Spearman and ANOVA were adjusted by Benjamini and Hochberg method[74].

### Elastic net models

Every tested model was built under 10-fold cross-validation using function cv.glmnet of R package glmnet (http://web.stanford.edu/∼hastie/glmnet/glmnet_alpha.html) with the following parameters: lambda.min.ratio=0.01 (the default) and nlambda=25 (default was 100). Parameter alpha varied as {0.1; 0.3; 0.5; 0.9; 1.0}. The reported cross-validated mean error and the number of variables in the model corresponded to lambda.1se, i.e. largest value of lambda found within 1 standard error of the minimum lambda. The regression of observed on predicted values was plotted using lambda.min.

### Drug sensitivity models in TCGA patients

We used the follow-up time profiles for which both status records “relapse/relapse-free” and “dead/alive” were available, which allowed creating “relapse-free survival” and “overall survival” variables. Depending on the cancer aggressiveness and chemotherapy type, different timeframes could become informative in the analysis of the eight TCGA cohorts. The follow-up timeframes were defined as 1/5^th^, 1/2^nd^, and full available (up to 18 years) intervals.

For the analysis reported in “Statistical power to detect correlates of drug sensitivity”, we used 42 drugs which were applied to at least 10 patients in one of the eight cohorts. In Figure 3 we report fractions of adjusted p-values (FDR) from this analysis calculated by Benjamini and Hochberg. For the analysis of “agreement between in vitro screen and clinical data” we only considered 14 of the compounds, which were found in the *in vitro* sets. The p-values from this analysis were Bonferroni-adjusted in the cross-comparisons between the *in vitro* and clinical results.

Matching significance of the drug-feature correlations that had been detected in the cell-line *in vitro* screens required accounting for multiple clinical variables. Such phenotype covariates as well as drug treatment data were obtained from TCGA as biotab files via https://tcga-data.nci.nih.gov/tcgafiles/ftp_auth/distro_ftpusers/anonymous/tumor/*/bcr/biotab/clin/

In order to measure and probabilistically estimate these effects, we fitted Cox proportional hazards regression models for every feature versus drug combination. Using all covariates available a cohort (such as “age at diagnosis”, “year of diagnosis”, “race”, “gender”, “ethnicity”) could result in unrealistically complex models. We thus included only covariates most likely associated with the disease prognosis, such as tumor degree, pathological tumor stage, immunohistochemical statuses in BRCA, Gleason score in PRAD, Karnofsky score in GBM (Suppl. Table 4). Next, we reasoned that when the association “feature - drug response” truly exists, we should observe it specifically in the patients who did receive the drug in the given TCGA cohort. Our survival models of the form

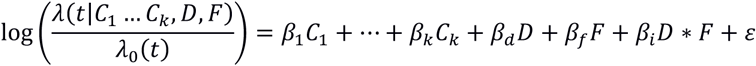

contained, apart from the covariates *C_1_…C_k_* and the residual term *ε*, main effects “drug” *D* and “feature” *F* as well as the interaction term *D*\**F*. A significant main effect of a drug could be interpreted as patients’ benefit in total and irrespective of the feature value, e.g. regardless of a gene mutation, or a gene expression, or a NEA-based pathway score. Conversely, a significant feature effect indicated that the feature correlated with survival directly, i.e. no matter if the drug was administered or not. Finally, significance of the interaction indicated efficacy of the drug specifically in patients with feature values either above or below a threshold, so that respective patterns could be explained by neither of the main effects. The interaction term was thus central for our purpose of detecting drug-feature correlations, whereas the significance of main effects of “feature” and “drug” was allowed although not required. As an example, a feature may or may not exhibit a significant correlation with survival in patients who did not receive the drug.

All survival analysis results were obtained using R package survival(http://dx.doi.org/10.1007/978-1-4757-3294-8). In order to estimate significance of the model terms, we used function coxph with continuous feature vectors. However, for visualizing the survival curves (Fig. 6) each feature was binarized at a cutoff that yielded the lowest p-value for the interaction term. Apart from the interaction model, we also checked if the p-value and FDR distributions preserved their properties under a unifactorial model. To this end, sub-cohorts of respective drug-treated patients were included in the survival analysis with the single main factor “feature”:

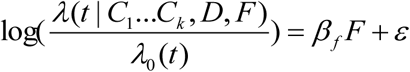

## ACKNOWLEDGEMENTS

The authors are grateful for help from National Bioinformatics Infrastructure Sweden (NBIS), the members of ACT (’Advanced Cancer Therapies’) consortium, and acknowledge financial support from Emil och Wera Kornells Stiftelse, Stockholm County Council, Swedish Cancer Society, Swedish Research Council, and Karolinska Institutet. We also recognize the contribution of specimen donors and research groups behind the samples of Cancer Cell Line Encyclopedia and The Cancer Genome Atlas.

## SUPPLEMENTARY FILES

| File | Description |
| --- | --- |
| SupplementaryTablesAndFigures.docx | Supplementary tables and figures |
| IC50values.ACTscreen.xlsx | IC50 values of drug sensitivity over the cancer cell lines in the ACT drug screen (see Methods). |
| glmnetModels.Basu_vs_new.raw.pdf | Building and validation of multivariate models of drug resistance from original point mutation and gene expression data.<br>As explained in Methods, the multivariate models were obtained using the elastic net algorithm under variable 'alpha' parameters (see values $A=0.1$; $A=0.3$; $A=0.5$; $A=0.9$; $A=1$ in the top left corners of each page). The algorithm tried to minimize the mean-squared error by reducing the number of features in the model (top legend in upper right plot). The final number of features as well as the 'lambda.1se' at which the practically best performance was achieved are indicated as 'N=' and 'L=' in the top left corner. The right and left vertical dotted lines show absolute minimum lambda and 'lambda.1se' found within 1 standard error of the former, respectively. The chosen features with their linear coefficients are listed below (sorted by coefficient values; the lists are truncated when too long).<br>The two bottom plots display model performance by matching drug sensitivity predicted for each cell line (X axes) on data used for training (blue points, left) and on newly obtained data from the ACT screen (green points, right). The model performance is measured with Spearman rank correlation between predicted and observed data points.<br>See also the legend to Figure 5. |
| glmnetModels.Basu_vs_new.pwnea.pdf | Building and validation of multivariate models of drug resistance from PWNEA scores obtained by using point mutation and gene expression data.<br>See the legend above. |

## List of abbreviations

AGS: altered gene set (gene set characterizing an individual sample/cell line/patient)
FGS: functional gene set (typically a pathway);
NEA: network enrichment analysis;
PWNEA: NEA at pathway level (i.e. by using multi-gene FGS);
GNEA: NEA by using single-gene FGS (i.e. individual network nodes);
ORA: overrepresentation analysis of FGS versus AGS;
GSEA: gene set enrichment analysis of FGS versus full ranked gene lists;
AGSEA: variant of GSEA where genes are ranked by absolute value;
ZGSEA: variant of GSEA where genes are ranked by z-score of deviation from cohort mean.

